# Role of a FAD-dependent monooxygenase in diazo group functionalisation of kinamycin in *Streptomyces ambofaciens*

**DOI:** 10.1101/2024.11.29.626053

**Authors:** Cláudia M Vicente, Alexis Boutilliat, Laurence Hôtel, Cédric Paris, Bertrand Aigle

## Abstract

Kinamycin biosynthesis is a complex process which has been extensively studied over the years, yet specific enzymatic steps continue to be unveiled. A diazo group present in the molecule is responsible for the promising antitumoral activity of kinamycins but its specific installation in *Streptomyces ambofaciens* is not fully understood. In this study, we explore the diazo functionalization of kinamycin in this strain. A new FAD-dependent monooxygenase is identified which is essential for kinamcyin biosynthesis. In its absence, stealthin C accumulates instead likely as a pathway shunt-product. Furthermore, as a result of the position of the gene encoding this monooxygenase, named *alp2F*, we also propose new boundaries of the kinamycin biosynthetic gene cluster resulting in a large cluster spanning over 72kb. This work paves the way for the continued understanding of the biosynthetic steps that are characteristic of diazo-containing natural products, and provides new biocatalysts for molecular engineering and accelerate bioactive compounds production.

## Introduction

Polyketide natural products constitute a chemically rich group of molecules often with important pharmaceutical and commercial applications. *Streptomyces*, Gram-positive bacteria with a high GC-content linear genome, are prolific producers and known to possess a large variety of biosynthetic gene clusters for natural products [1–3]. Such a vast solution space can hinder the discovery of new molecules as well as the characterization of specific biosynthetic steps of previously identified compounds [4–6]. The diazo group constitutes a type of N–N bond which has been identified in more than 300 natural products in the past few decades (Figure 1) [7, 8]. Diazofluorene polyketides such as kinamycin and lomaiviticin have diazo functionalities integrated into a benzo[*b*]fluorene scaffold [9]. This structural feature endows kinamycin and lomaiviticin with promising DNA-related cytotoxic and antitumor activities [10–12]. To date, only four *Streptomyces* species have been described to possess the kinamycin biosynthetic cluster and to produce it, namely *Streptomyces ambofaciens* [13, 14], *Streptomyces murayamaensis* [15], *Streptomyces chattanoogensis* [16] and *Streptomyces galtieri* [17]. Like most polyketides, kinamycin has a complex biosynthetic pathway. It proceeds via the dehydrorabelomycin, that is produced from the condensation of one acetyl-CoA and nine molecules of malonyl-CoA through the action of PKSs and oxidases. Then, a ring contraction followed by other modifications occur to obtain prekinamycin, the first pathway intermediary to contain the diazo group. Prekinamycin is then converted into the final product kinamycin. In *S. ambofaciens* the genes encoding the enzymes responsible for kinamycin biosynthesis are organized in the *alp* cluster [13, 14]. Since its discovery in the early 70s, extensive work has been carried out to elucidate the kinamycin biosynthetic pathway [18–20, 14]. As it was initially described, the *alp* cluster contained 28 genes including at least five putative oxidases. These modifying enzymes are responsible for post-PKS specific steps in the biosynthesis. For example, oxidases AlpK and AlpJ are involved in the contraction of the B-ring [21]. However, often the precise role of post-PKS genes is not fully elucidated. Diazo groups have been shown to be key for the remarkably enhanced cytotoxicity found in both kinamycin and lomaiviticin [22]. The study of diazo assembling first made the most progress focusing on another antibiotic, cremeomycin from *Streptomyces cremeus*. Although not a polyketide like kinamycin, cremeomycin has a diazoketone structure and two genes have been described to be involved in the synthesis of the diazo group. These are *creE* encoding a FAD-dependent monooxygenase, and *creD* which encodes a lyase [23, 24]. In the biosynthetic pathway of cremeomycin, the FAD-dependent monooxygenase is proposed to carry out sequential oxidations to form nitrosuccinic acid from L-aspartic acid, while the lyase mediates the β-elimination of nitrosuccinic acid to form nitrous acid that is introduced in the final molecule to give cremeomycin [24]. A study using heterologous expression to reconstruct the kinamycin biosynthetic pathway from the strain S. *galtieri* proposed stealthin C as an intermediary [17]. More recently, Wang et al. studied fosfazinomycin and kinamcyin biosynthesis from *Streptomyces* sp. NRRL S-149 and S. murayamaensis ATCC21414 and demonstrated that diazo assembly actually proceeds independently and is then transferred into the scaffold using glutamic acid as a carrier [25, 26]. Despite these breakthroughs, thus far no enzymes related to the synthesis of the kinamycin’s diazo group specifically in the *alp* cluster from *S. ambofaciens* have been characterized. Here we identify a new FAD-dependent monooxygenase, investigate its role in the diazo assembly in the kinamycin biosynthetic pathway of *S. ambofaciens* and assign it to the *alp* cluster. Elucidating the formation of these unique chemical moieties is important not only to achieve a better understanding of kinamycin biosynthesis but also supports the development of structural diversity in other bioactive polyketides.

**Figure 1.**
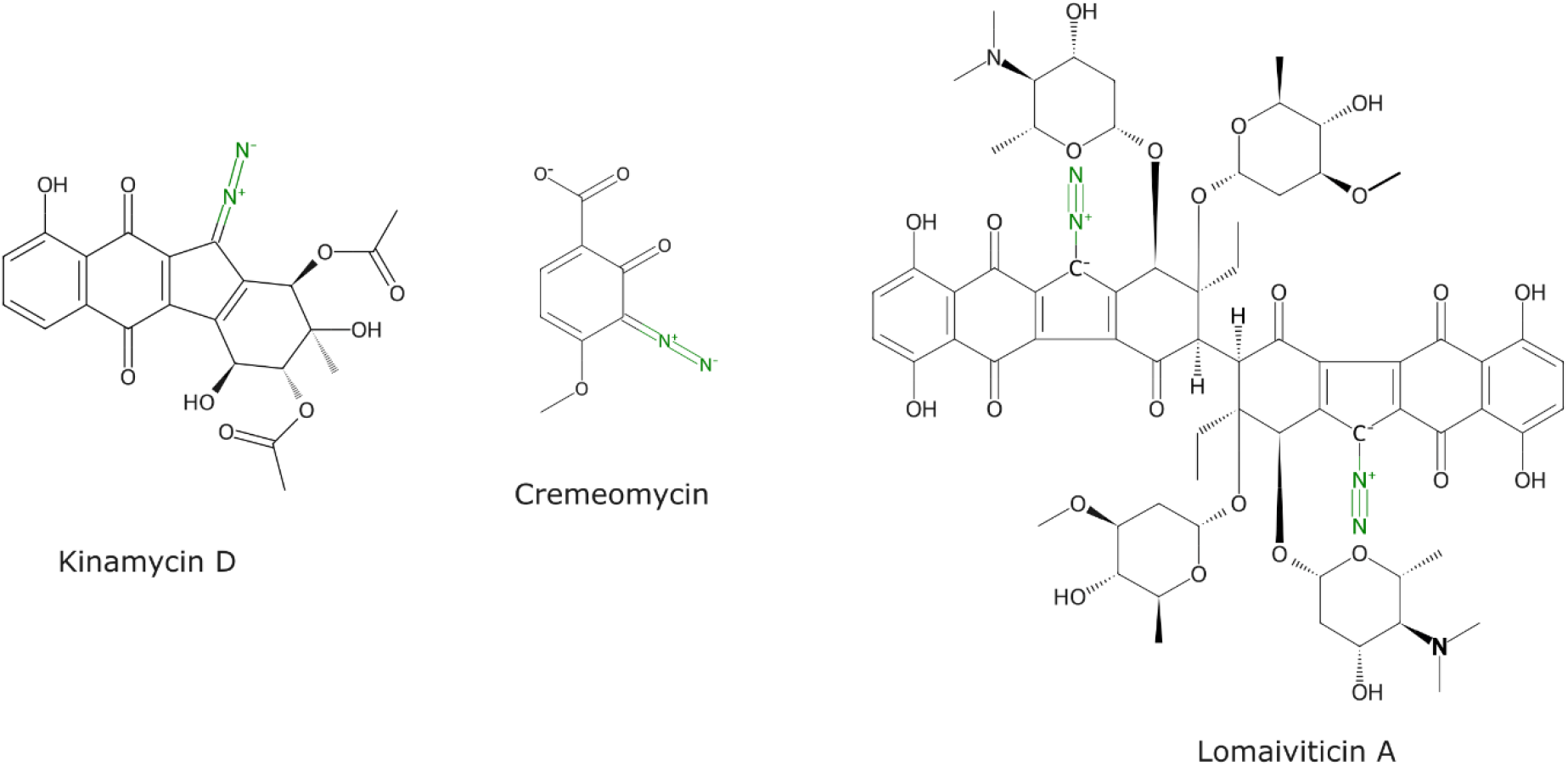
Selected natural products containing diazo groups.

## Materials and Methods

### Strains, media and growth conditions

Strains and plasmids used in this work are listed in Table S1. *Streptomyces* strains were routinely grown in TSB and R2 media at 30ºC, while sporulation was carried out in MS medium also at 30ºC [27]. LB medium was used for *Escherichia coli* and *Bacillus subtilis* cultures at 37 and 30ºC, respectively [28]. Unless indicated otherwise, solid versions of the media were prepared with 2% agar. Antibiotics apramycin, kanamycin (50 µg/ml), chloramphenicol and nalidixic acid (25 µg/ml) were added when required. All strains were maintained as spores (*S. ambofaciens* and *B. subtilis* strains) or cell suspensions (*E. coli* strains) in 20% (v/v) glycerol and stored at -80ºC.

For the fermentation of *S. ambofaciens* ATCC23877 and its derived strains, serial dilutions of the stock were spread on fast-growth solid medium TSB and the plates incubated 24h at 30ºC for viable spores’ quantification. Cultures were performed in 50 ml R2 medium in 250 ml shake flasks inoculated with 5x10^6^ viable spores and incubated at 250 rpm and 30ºC until antibiotic production onset at late exponential-stationary transition phase.

### Sequence analysis

The DNA sequence of *creE* from the cremeomycin producer *S. cremeus* NBRC 12760 was used to search homolog genes in the genome of *S. ambofaciens* using BLAST [29]. The tools cblaster and clinker were used to search homologs in other *Streptomyces* biosynthetic gene clusters and to visualize the aligned clusters [30, 31].

### DNA manipulations and materials

*Streptomyces* genomic DNA was isolated as described previously [27]. *E. coli* DNA manipulations, competent cells preparation and transformation were performed as described previously [28].

### Strain engineering

An *in-frame* deletion of both copies of the gene *alp2F*, each located in one of the chromosomal arms (SAM23877_0158 and SAM23877_7516, accession numbers AKZ53207.1 and AKZ60557.1, respectively) was made in the wild-type *S. ambofaciens* ATCC23877 strain to create the ΔΔ*alp2F* mutant strain. The strategy used was based on the PCR-targeting-based system REDIRECT [32]. Briefly, an *aac(3)IV*-oriT marker flanked by the same regions as the target gene *alp2F* was obtained by PCR and used to first replace the gene in the library cosmid E8 through homology recombination in the strain *E. coli* BW25113/pIJ790. The resulting E8Δ*alp2F* mutant cosmid was then transferred into the *S. ambofaciens* wild-type strain through interspecific conjugation [27]. Exconjugants with both genomic gene copies replaced by the resistance marker (double-crossover mutants) were selected through the correct selection marker resistance pattern (apra^R^, kan^S^). Two independent mutants were verified by PCR using flanking primers to the insertion region (Table S2), sequencing and southern blot analysis (data not shown).

### Complementation of the mutant strain

To validate the functional characterization of *alp2F*, the gene mutation was complemented with the *alp2F*-*alp2G* operon using the ϕBT1-derived integrative plasmid pRT802 [33]. The entire operon, including its predicted promoter and terminator regions (4.1kb size insert), was first amplified using the primers listed in Table S2 and digested with *EcoR*I/*BamH*I for the insertion into pRT802 digested with the same enzymes. Cloning was verified through PCR amplification and sequencing (data not shown). The complemented strain was obtained transferring this construction pRT802::*alp2F-alp2G* into two independent clones of the mutant strain ΔΔ*alp2F* through interspecific conjugation. Exconjugants harbouring the plasmid were selected through antibiotic resistance (apra^R^, kan^R^) and two independent complemented clones were further confirmed through colony PCR (data not shown).

### Metabolite extraction

Antibiotic production was assessed on R2 medium as described previously for each of the independent clones [13, 19]. Samples were collected upon antibiotic production onset, then quenched and extracted with 1 volume of 99% ethyl acetate and 1% acetic acid with shaking at 150 rpm for 1h at 4ºC. The organic phase was then dried in a rotavapor (Buchi R-II, Switzerland) and the resulting extracts were dissolved in methanol 50% in a 1/1000 of the initial extraction volume.

### Kinamycin bioassays

The presence of kinamycin was assessed through bioassays using *B. subtilis* ATCC6633 as an indicator strain. In brief, a soft LB plate (0.6% agar) was prepared mixing the medium with 10^6^ viable spores of the indicator strain. Then, 5 µl of the crude extracts were deposited onto a paper disk previously placed on the plate. Solutions were let to diffuse for 1h at 4ºC, followed by incubation at 30ºC o/n.

### Liquid chromatography-mass spectrometry (LC-MS)

Qualitative and semi-quantitative analyses of metabolites were performed by LC-HRMS on a UHPLC-MS system (Thermo Fisher Scientific, San Jose, CA, USA) consisting in a quaternary UltiMate3000^TM^ solvent delivery pump connected to a photodiode array detector (PDA) and an LTQ-Orbitrap^TM^ hybrid mass spectrometer.

Sixteen microliters of crude extracts prepared as described previously, were separated on a C18 Alltima reverse phase column (150 x 2.1 mm, 5 µm – Grace/Alltech, Darmstadt, Germany) equipped with a C18 Alltima pre-column (7.5 x 2.1 mm, 5 µm) at 25°C. The flow rate was set at 0.2 ml.min^-1^ and mobile phases consisted in water modified with formic acid (0.1%) for A and acetonitrile modified with formic acid (0.1%) for B. Compounds of interest were separated using the following elution profile: 0– 48 min, linear gradient 5–95% solvent B; 48–54 min, constant 95% solvent B; 54–54.1 min, linear gradient 95-5% solvent B, 54.1-60 constant 5% solvent B.

Mass analysis was carried out in ESI positive ion mode (ESI+) and mass spectrometry conditions were as follows: spray voltage was set at 5.0 kV; source gases were set (in arbitrary units.min^-1^) for sheath gas, auxiliary gas and sweep gas at 30, 10 and 10, respectively; capillary temperature was set at 275°C; capillary voltage at 4 V; tube lens, split lens and front lens voltages at 155 V, - 28 V and - 6.00 V, respectively. Ion optics parameters used were previously optimized by automatic tuning using a solution of stambomycin (0.1 g/l), another polyketide produced by *S. ambofaciens*.

Full scan MS spectra were performed at high resolution (R=60000) on the Orbitrap^TM^ analyser from 120 to 2000 m/z to obtain exact masses of metabolites. MS2 full scan spectra were additionally realized for structural elucidation thanks to LTQ^TM^ analyser (Linear Trap Quadrupole). Raw data were processed using the XCALIBUR^TM^ software program (version 2.1, Thermo Fisher Scientific).

## Results

### FAD-dependent monooxygenase encoding gene identified within the *alp* cluster

The genome of *S. ambofaciens* ATCC23877 (accession number CP012382.1) was searched to identify genes encoding for FAD-dependent monooxygenases based on the sequence of *creE* from the cremeomycin producer *S. cremeus* NBRC 12760. The results revealed the presence of two *creE*-like genes (SAM23877_0158 and SAM23877_7516) in the neighbouring regions of the two copies of the a*lp* gene clusters, located in the terminal inverted repeated regions and duplicated as the *alp* cluster (Figure 2). Further analysis showed that SAM23877_0158 (or SAM23877_7516) encodes a monooxygenase from the FAD/NAD-Binding Rossmann superfamily (pfam13454), with an 60.4% identity with CreE and which belongs to the same protein superfamily, suggesting both proteins may share enzymatic functions. A search for protein sequences available on genome databases revealed that homologues of SAM23877_0158 are widely distributed in the *Streptomyces* genus, including producers of relevant natural products. These include *S. ambofaciens* DSM 40697, a phylogenetically-close strain to *S. ambofaciens* ATCC23877 and producer of kinamycin (99.7% identity) [34], *S. glaucescens* strain GLA.O (63.1% identity), *S. cattleya* NRRL 8057 (63.1% identity), the *S. bingchenggensis* BCW-1 strain (56.4% identity), and *S. davaonensis* JCM 4913 (60.2% identity), among others (Figure S1).

**Figure 2.**
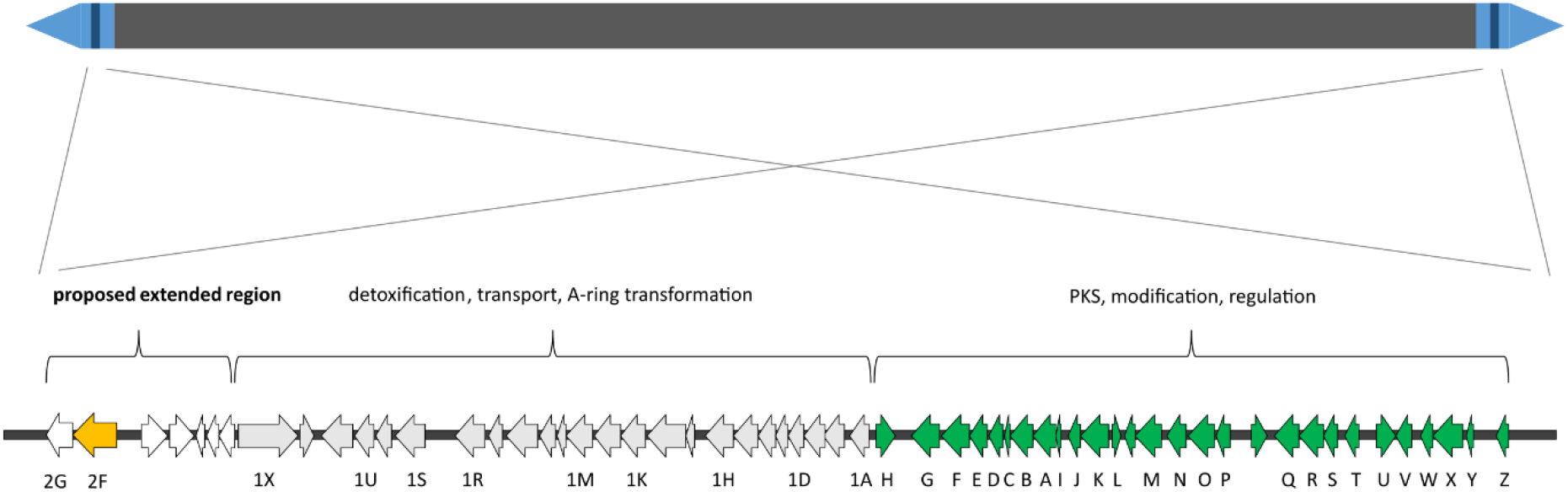
Localization and organization of the kinamycin biosynthetic cluster. The terminal inverted repeats are represented in light blue, the *alp* cluster in dark blue bars. The genes belonging to the first described alp cluster which encoded the PKS and modification enzymes as well as the transcriptional regulators are in green. The genes involved in detoxification, transport and A-ring transformation are in grey. The FAD-dependent monooxygenase encoding gene *alp2F* here described is highlighted in yellow and the proposed extended region is indicated.

The gene SAM23877_0158 was found to be upstream of a lyase encoding gene, SAM23877_0157, that is also duplicated (SAM23877_7515) and shows an 67.0% identify with CreD from *S. cremeus*. Surprisingly, this genetic organization is highly conserved among streptomycetes and can be found in all the strains previously identified with homologs of SAM23877_0158 (Figure S1).

### Monooxygenase activity is essential for kinamycin biosynthesis

Due to its location in a region adjacent to the *alp* cluster and its predicted function, SAM23877_0158 could potentially be involved in kinamycin biosynthesis. To test this hypothesis, a mutant strain was engineered with both copies of the gene deleted (i.e., SAM23877_0158 and SAM23877_7516). When the wild-type and mutant strains were grown on solid media in producing conditions, there was a clear metabolic difference. While the wild-type strain plate showed the characteristically dark orange pigment that is associated with kinamycin, the plate with the mutant strain lacking SAM23877_0158 was distinctly pink, suggesting the absence of this compound and/or the presence of another metabolite (Figure 3). In order to evaluate kinamycin biosynthesis, fermentations of both strains were performed in liquid media, and the obtained crude extracts were analysed using bioassays. It is worth mentioning that in the selected conditions none of the other antibiotics produced by *S. ambofaciens* are detected [13, 14]. Upon observation, the extracts presented the same colours as previously observed on plate (Figure S2). In the bioassays the mutant strain pink extract showed no antibiotic properties, in contrast with the orange extract, indicating that kinamycin production is likely abolished in the mutant strain (Figure 4). The two independently obtained clones of the mutant strain presented the same phenotype, confirming the mutation leads to the absence of kinamycin in the samples. To further validate gene function, we decided to complement the mutation introducing a conjugative-integrative plasmid carrying a copy of the operon SAM23877_0157-SAM23877_0158 under its own promoter. Bioassays were performed with crude extracts of the complemented strain. When the empty plasmid was introduced in the wild-type or in the mutant strain, no change in kinamycin production was observed compared to the strains without the plasmid (data not shown). However, when a copy of the operon SAM23877_0157-SAM23877_0158 was introduced in the mutant to yield the complemented strain, the resulting extracts regained the same orange colour as in the wild-type strain extracts (Figure S2). Moreover, the bioassays showed that kinamycin production is also restored to levels similar to those of the wild-type strain (Figure 4). Taken together, these results confirm the important role of this FAD-dependent monooxygenase in kinamycin biosynthesis. They also suggest that despite its localization in what was initially thought to be an adjacent region to the *alp* gene cluster, SAM23877_0158 effectively belongs to the kinamycin biosynthetic gene cluster. It has hence been renamed *alp2F* ensuing the gene order described in the most recent publications featuring the *alp* cluster [17]. Likewise, the adjacent gene SAM23877_0157, which is predicted to belong to the same transcriptional unit, is named *alp2G* (Figure 2).

**Figure 3.**
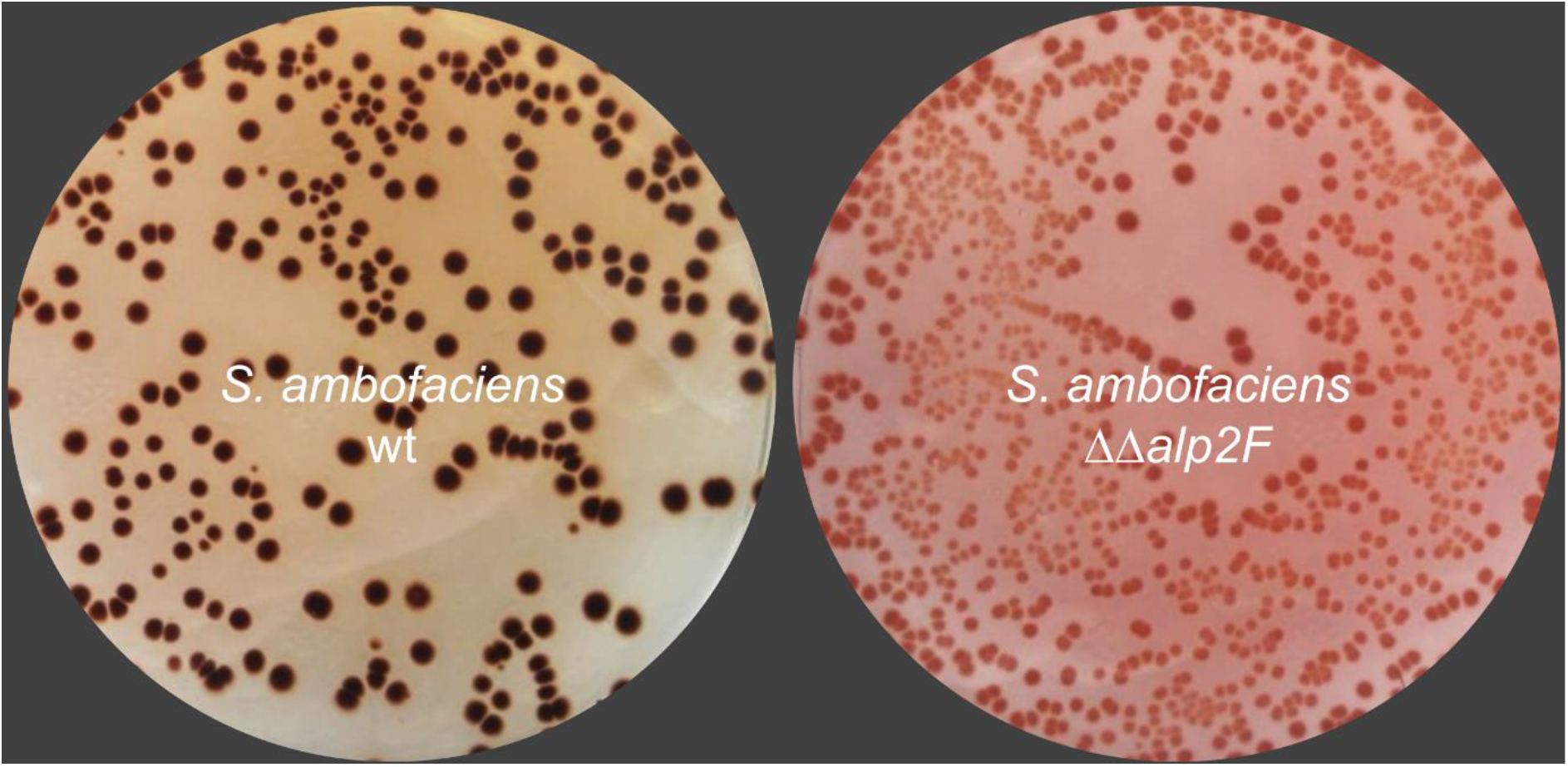
Comparison of *S. ambofaciens* wild-type and mutant strains. Kinamycin producing conditions in solid media were used, resulting in dark orange colonies and surrounding media in the wild-type strain, that contrasted with the pink coloration in the mutant strain where both copies of *alp2F* are deleted.

**Figure 4.**
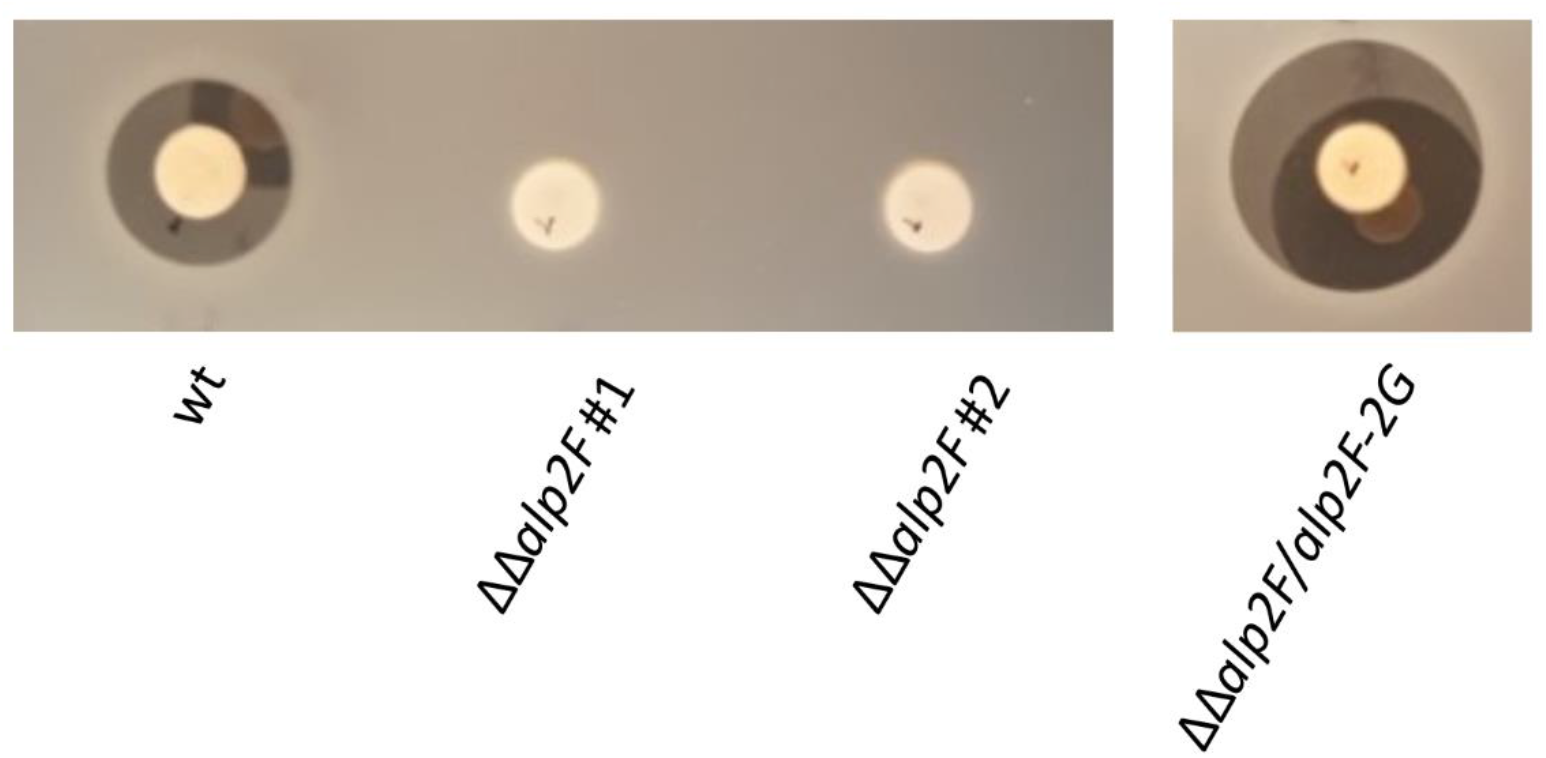
Antibiotic assay for kinamycin detection. The mutation of *alp2F* leads to the loss of kinamycin production, as can be observed comparing the *S. ambofaciens* wt strain with two independently obtained clones of the mutant strain ΔΔ*alp2F*. The complemented strain ΔΔ*alp2F/alp2F-alp2G* has kinamycin production restored. The culture and extraction conditions used in this assay ensure that no other antibiotic activity is detected.

### Stealthin C is a kinamycin biosynthesis derived metabolite

The biosynthetic pathway of kinamycin involves multiple steps carried out by several different enzymes. The characteristic pink colour of colonies and extracts from the ΔΔ*alp2F* mutant strain suggested the presence of an unknown metabolite, albeit without antibiotic activity in the tested conditions. We postulated that kinamycin, the final product of the biosynthetic pathway, is not being synthesized due to a missing enzymatic reaction. Instead, the pathway stops at this unknown intermediate metabolite. An LC-MS analysis was therefore performed to identify the compounds present on the crude extracts. Results showed that kinamycin D (monoprotonated form with m/z 455.1085) was only present in the sample from the wild-type strain (Figure 5A, top panel). The ΔΔ*alp2F* mutant strain that lost the kinamycin producing capability instead accumulated another metabolite (Figure 5A and B, middle panels). This compound was barely present in the wild-type strain, suggesting it was either produced in very low amounts or was nearly completely transformed into the next intermediary of the pathway (Figure 5B, top panel). From the structural features of the kinamycin molecule and its predicted biosynthetic pathway, we hypothesised that this compound could correspond to stealthin C (C_18_H_13_NO_4_, M_mono_ = 307.0845). The newly detected peak in ΔΔ*alp2F* showed a parent ion with m/z = 308.0917 that matched perfectly the monoprotonated form of stealthin C expected at m/z_theo_ = 308.0918 (Figure 5B, middle panels). Moreover, the UV profile of this peak showed the same absorption maxima as those previously described for stealthin C (Figure S3) [36]. Taken together these results might suggest that the kinamycin biosynthetic pathway includes a single nitrogen atom containing intermediary or shunt product before the synthesis of prekinamycin. Other pathway derived metabolites were also searched, such as seongomycin ([M+H]^+^ form expected at m/z = 454.0960) which was detected in all the strains, whereas intermediary kinobscurinone ( [M+H]^+^ form expected at m/z = 306.0528) was not detected in any of the samples likely due to its low stability (Figure S4). As expected, kinamycin D was again found in the extracts when Alp2F is present in the complemented strain ΔΔ*alp2F*/*alp2F-alp2G*, as well as some residual stealthin C (Figure 5, bottom panel).

**Figure 5.**
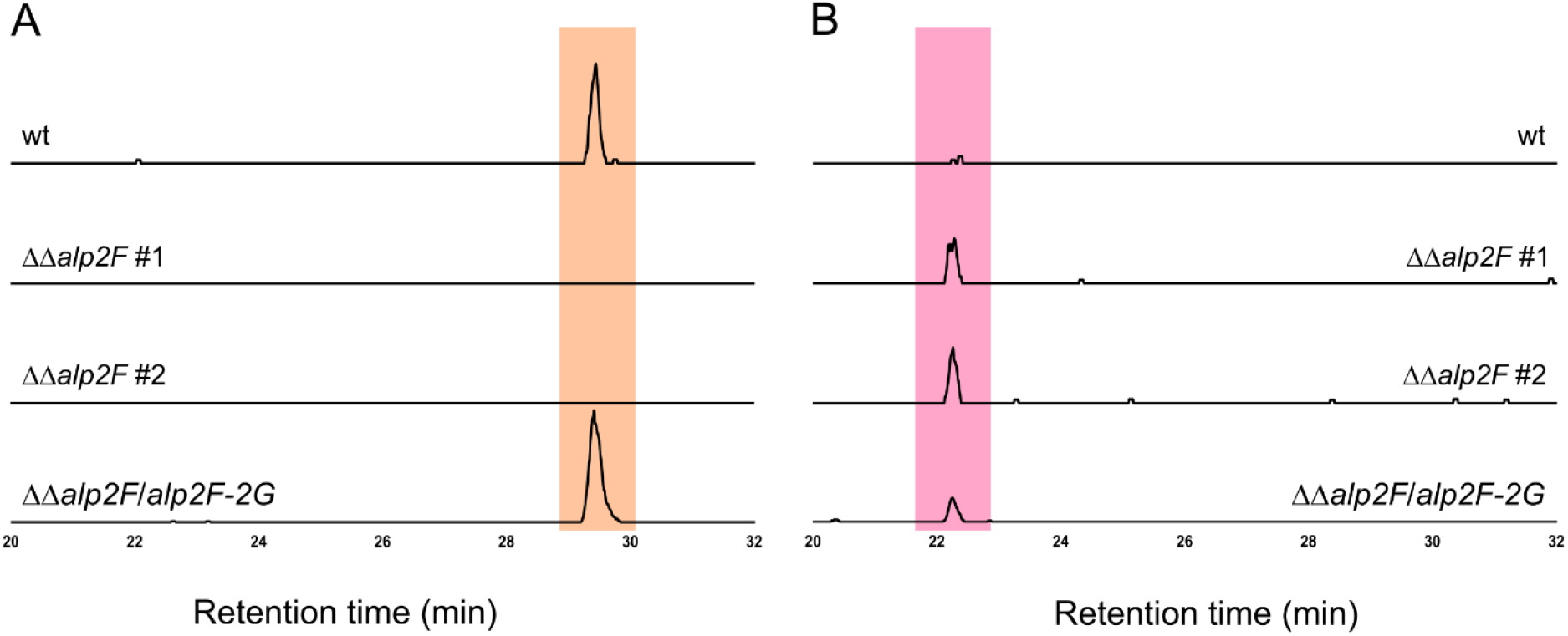
LC-MS analysis of the metabolites produced by the different strains. Detection of kinamycin D was set at m/z [M+H]^+^ 455.1085 (panel A, highlighted in orange), stealthin C at m/z [M+H]^+^ 308.0917 (panel B in pink). Analysis was performed on crude extracts prepared from the wild-type strain, two independent clones of the mutant strain ΔΔ*alp2F* and from the complemented strain ΔΔ*alp2F/alp2F-2G*.

### Proposed new *alp* cluster boundaries

The discovery of the new gene *alp2F* encoding a FAD-dependent monooxygenase that participates in the kinamycin biosynthetic pathway puts into question the size of the currently established *alp* cluster in *S. ambofaciens*. We therefore propose to change the boundaries of the *alp* cluster to include *alp2F* as well as the other gene in the operon *alp2G*. The newly suggested cluster contains a total of 59 genes ranging from *alp2G* next to the here described *alp2F* gene until *alpZ* (Figure 2). It is well recognized that genes involved in the same process are typically clustered together. Although further studies are required to test this hypothesis, it is likely that the five genes located between *alp2F* and *alp1X*, belong to the same cluster and play a role in kinamycin biosynthesis.

## Discussion

Diazo groups have long been the subject of interest in synthetic and biological chemistry because of their exceptional reactivity. This N-N bond is directly responsible for the promising cytotoxic activity of kinamycin [12, 22]. In addition to kinamycin, only six other biosynthetic clusters have been identified for diazo-containing compounds, namely cremeomycin, alazopeptin, tasikamide, avenalumic acid, azaserine and lomaiviticin [24, 37–41]. The formation of this structural feature has remained elusive, particularly in the case of kinamycin biosynthesis. It has been demonstrated that stealthin C can be obtained from L-cysteine through the action of AlpJ, another FAD-dependent monooxygenase [36, 42]. The same work also questioned the role of stealthin C in the pathway and for the first time suggested that it can be obtained non-enzymatically. A recent study focused on the *alp* cluster from *S. galtieri* and used heterologous expression in the more genetically tractable *S. albus* to reconstruct the biosynthetic pathway. Genes *creE* and *creD*-like, homologous to *alp2F* and *alp2G* respectively, were identified and their involvement in the biosynthesis was confirmed with deletion mutants that lost the ability to produce kinamycin D [17]. However, no accession number was given for the gene sequences and no other metabolites were identified in these mutant strains, providing no further clues on their roles.

Here, a new FAD-dependent monooxygenase encoding gene was identified just on the outskirts of the *alp* cluster in *S. ambofaciens*, named *alp2F*. We demonstrate through functional characterization and metabolite identification that Alp2F participates in kinamycin biosynthesis. In the absence of *alp2F*, kinamycin D production is abolished and a new metabolite accumulates instead. A high-resolution mass screening (tolerance of 10ppm) was used to confidently identify it as stealthin C. The fact that no obvious daughter ions were observed through MS/MS experiments is coherent with the molecular structure of stealthin C, further validating its identification. Moreover, the characteristic pink colour of stealthin C has been reported in early studies [43]. Although stealthin C was initially described as an intermediary of the kinamycin biosynthetic pathway and the target of the diazo installation [44] in a process similar to the diazo formation in cremeomycin [23], this hypothesis has been steadily losing support. Despite the accumulation of stealthin C in high levels, we propose it is a shunt-product of kinamycin biosynthesis as the result of a non-enzymatic reaction from kinobscurinone (Figure 6). Based on the confirmed intermediates that were detected, we hypothesise that diazo installation passes through a separate molecular entity and *alp2F* is involved in the synthesis of this N-N carrier rather than on stealthin C itself (Figure 6). This is in good accordance with recent studies that identified glutamylhydrazine as the hydrazine donor in both kinamycin and fosfazinomycin biosynthesis [25]. Moreover, the biosynthetic enzymes FzmR, FzmQ, FzmO and FzmN involved in the synthesis of glutamylhydrazine have homologs in the *alp* cluster, Alp1N, Alp1M, Alp1L and Alp1K, respectively, which coincidently have previously been proposed to be the part of the diazo assembly machinery [45]. The long-time known *alpH* gene has recently been described as encoding a O-methyltransferase-like enzyme which catalyses a SAM-independent installation of glutamylhydrazine onto the polyketide scaffold during kinamycin biosynthesis [26], putting to rest the discussion around stealthin C and definitely confirming its status as a shunt-product in *S. ambofaciens*.

**Figure 6.**
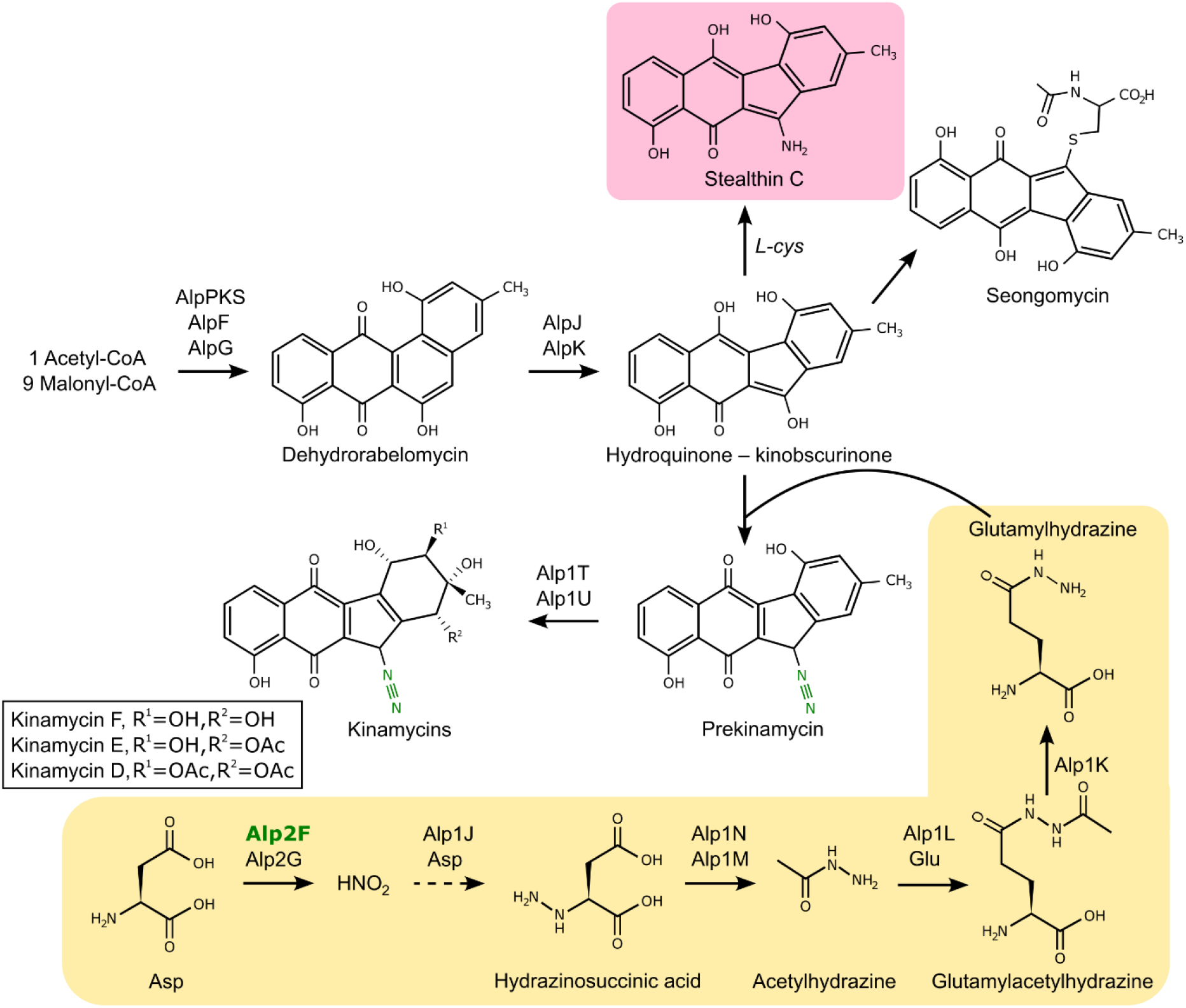
Kinamycin biosynthetic pathway. The proposed pathway includes the shunt metabolite stealthin C and the role of Alp2F in the formation of the diazo group, based on the model proposed in [25].

Expanding our knowledge on the different enzymes involved in kinamycin biosynthesis such as *alp2F* can potentially have implications in natural products synthetic biology. Homologous genes to *creE* and *creD* identified in *S. davawensis* have been demonstrated to lead to the synthesis of alternative derived molecules [46]. From a purely fundamental point of view, these results add another gene to the *alp* cluster. The fact that the first studies on *S. cremeus* did not identified homologs in the kinamycin cluster from *S. ambofaciens* [24] probably comes down to the cluster delimitations as they were described at the time. Here, we demonstrate that *alp2F* should be included in the kinamycin biosynthetic cluster, supporting a revision of its boundaries. This is not the first time that the *alp* cluster’s limits are modified. Since its first description with 29 genes from *alpH*-*alpZ*, an adjacent region containing 24 genes from *alp1X*-*alp1A* and located next to *alpH* was added to the cluster [21]. In a more recent study, an *afsA*-like gene involved in the biosynthesis of a signalling molecule and located in the other extremity after *alpZ* was proposed to also belong to the alp cluster [47]. The predicted *alp2G* located downstream of *alp2F* almost certainly belongs to the cluster, although it was not the object of this work and further studies are necessary to investigate its function. We hypothesise that this lyase-encoding gene is involved in the production of nitrous acid from asparagine, in a similar fashion as *creE* does it from nitrosuccinic acid in the cremeomycin biosynthesis [24]. Adding the *alp2F* gene results in a cluster with a total of 59 genes (Figure 1). *Streptomyces* are known to possess a variety in biosynthetic gene clusters that can have a considerable size and occasionally reach over 100kb. In *S. ambofaciens*, the *alp* cluster spans therefore over 72kb (over 87 kb if included the afsA-like gene), making it the strain’s third largest gene cluster after the spiramycin and stambomycin gene clusters (with 90kb and 146kb, respectively) [47].

Overall, this study sheds another light on diazo group assembly in kinamycin as well as in the variety of very specific oxidases that can be present in a single gene cluster, further demonstrating that our knowledge of the complexity of these biosynthetic pathways is still incomplete. Elucidating the installation of moieties such as diazo groups can provide new insights on compound modifications and lead to the engineering of synthetic metabolic pathways for alternative natural products with improved biological activities.

## Author statements

### Supplementary Material

Supplemental material, including four figures and two tables, is available online.

## Author contributions

CMV designed the study, performed the experiments, analysed the data, wrote the original draft and reviewed and edited the manuscript. AB and LH performed the experiments CP carried out the LC-MS analysis and spectra breakdown. BA designed the study, analysed the data, reviewed and edited the manuscript, acquired the funding and directed the research. All authors have read and approved the final manuscript.

## Conflicts of interest

The authors declare that there are no conflicts of interest.

## Funding information

This work was funded by the French National Research Agency (ANR) through a part of the “Investissements d’Avenir” program (ANR-11-LABX-0002-01, Lab of Excellence ARBRE) and through MiGenis project (ANR-13-BSV6-0009). CMV was also supported by the AgreenSkillsPlus Program (FP7609398.0000) and the Région Grand Est.

## Acknowledgements

LC-MS analysis was performed at the Structural Metabolomics Analyses Platform (PASM, SF4242, Université de Lorraine, France). The authors thank Christophe Corre (University of Warwick, UK) for critical reading of the manuscript and helpful suggestions.

## Notes

### Competing Interest Statement

The authors have declared no competing interest.

