## Supplementary material for "Role of a FAD-dependent monooxygenase in diazo group functionalisation of kinamycin in *Streptomyces ambofaciens*"

**Table S1**. Strains and plasmids used in this study.

| Strains / plasmids | Relevant properties^a^ | Source |
| --- | --- | --- |
| *S. ambofaciens* ATCC23877 | wild-type strain | [1] |
| *S. ambofaciens* ΔΔ*alp2F* | SAM23877_0158 locus replaced by the *aac(3)IV*-*oriT* cassette | This study |
| *S. ambofaciens* ΔΔ*alp2F*  /pRT802:: *alp2F-alp2G* | mutant strain complemented with *alp2F* | This study |
| *E. coli* |  |  |
| DH5α | cloning strain | [2] |
| BW25113 | strain used for gene replacement/mutation | [3] |
| ET12567/pUZ8002 | donor strain in interspecific conjugation | [4, 5] |
| *B. subtilis* ATCC6633 | indicator strain for bioassay antibiotic detection |  |
| *Plasmids/cosmids* |  |  |
| E8 | *S. ambofaciens* ATCC23877 genomic library cosmid | [6] |
| pIJ773 | *oriT*, *aac(3)IV* | [7] |
| pIJ790 | *oriT*, *exo*, *cat, gam*, *bet* | [7] |
| E8Δ*alp2F*::*aac(3)IV* | *alp2F* is replaced by *aac(3)IV*-*oriT* | This study |
| pRT802 | φBT1-integrative plasmid, *neo*, *attP* | [8] |
| pRT802::*alp2F-alp2G* | pRT802-derived plasmid with the SAM23877_0158-SAM23877_0157 locus | This study |

*aac(3)IV*, apramycin resistance gene; *cat*, chloramphenicol resistance gene; *neo*, kanamycin resistance gene; *gam*, inhibits host exonuclease V; *bet*, single-stranded DNA binding protein; *attP*, attachment site of φBT1; *oriT*, origin of transfer.

**Table S2.** Primers used in this study.

| Name | Sequence (5’-3’) | Function |
| --- | --- | --- |
| amont_creE | gccgccgagcacaggaacatggcgggcaccgggcacatgATTC  CGGGGATCCGTCGACC | selection marker amplification for mutant construction |
| aval_creE | ggcaggcggggccggcgggggcgggctggtcgcggctcaTGTA  GGCTGGAGCTGCTTC |  |
| creED_amont | CATAggatccCCCTCCTCCTCCTGGACG | mutation verification and locus amplification for plasmid construction |
| creED_aval | ACAGgaattcGGTCTGAGCAACTTGTGGC |  |

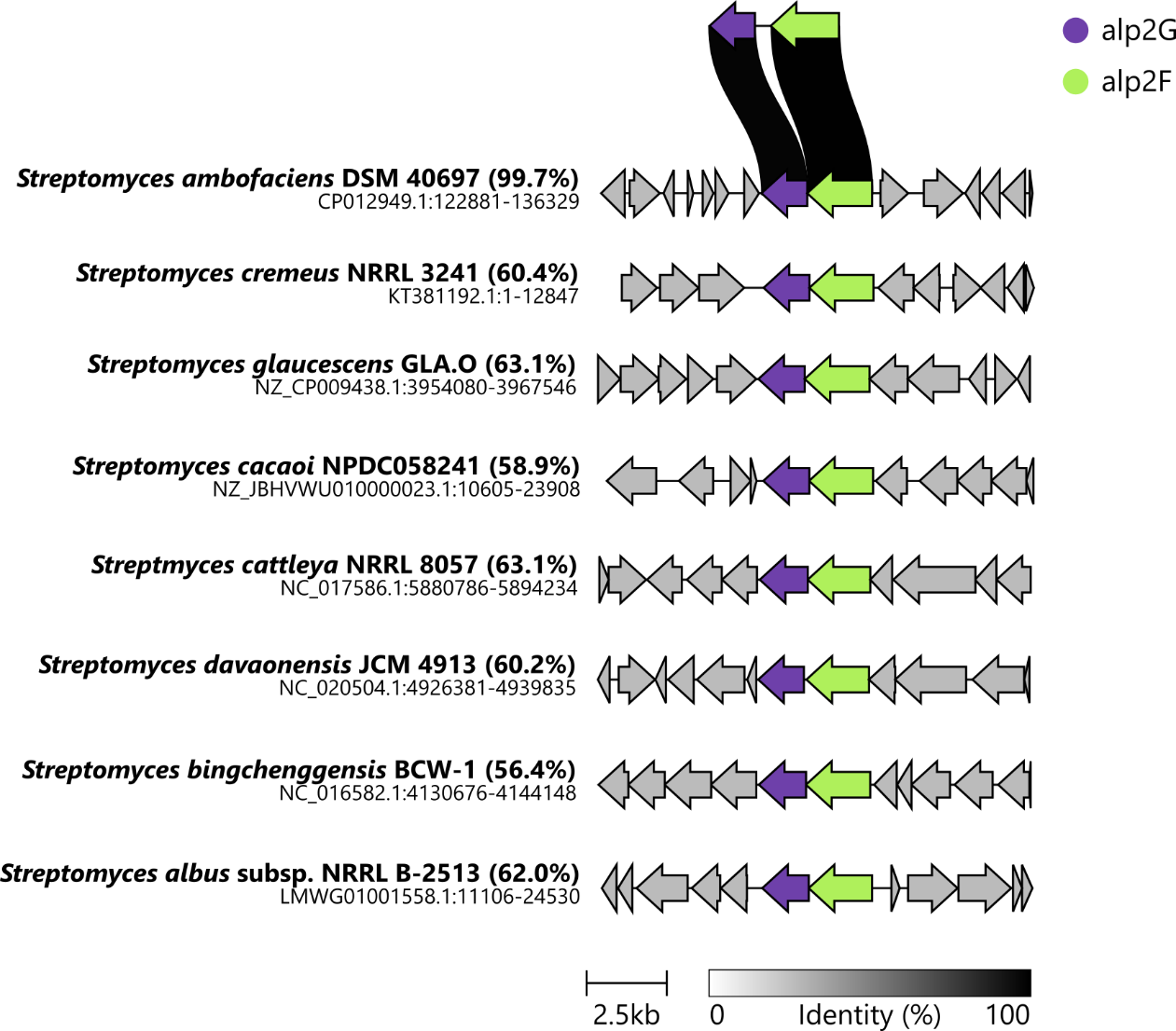

**Figure S1. Alignment of clusters containing alp2F and alp2G-like genes found in Streptomyces.** Identify comparing alp2G and alp2F from S. ambofaciens ATCC23877 with the other strains.

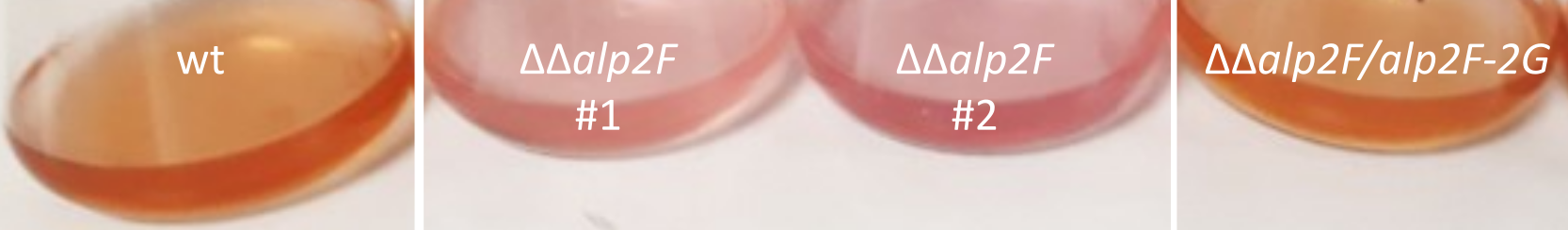

**Figure S2. The presence of alp2F gene results in differently coloured fermentation extracts.** The crude extracts obtained using an acidified extraction solution from the mutant strains ΔΔalp2F (99% ethyl acetate and 1% acetic acid; see Materials and Methods) present a pink colour compared to the orange wild-type extracts. The orange coloration phenotype is restored when the gene is reintroduced in the complemented strain ΔΔalp2F/alp2F-alp2G.

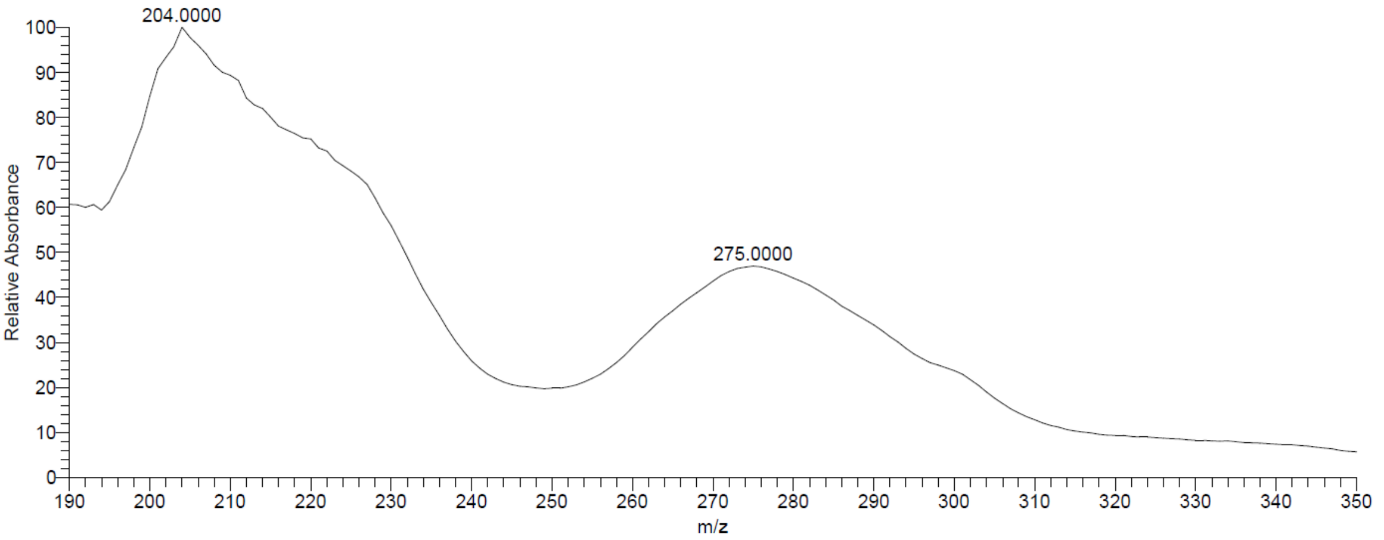

**Figure S3.** **UV absorption spectrum of peak identified as stealthin C.**

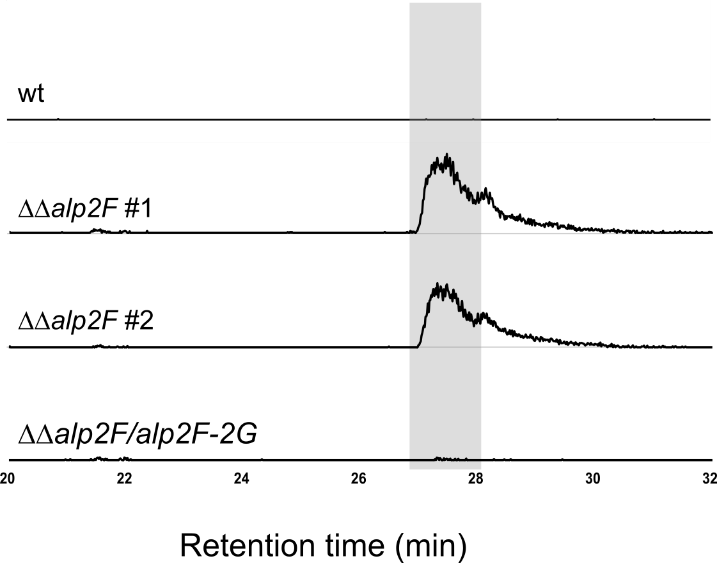

**Figure S4.** **LC-MS analysis of other metabolites in the kinamycin biosynthetic pathway in the different strains.** Seongomycin at m/z=454.0960 under its monoprotonated form (highlighted in gray) was detected in the mutant strain ΔΔalp2F but not in the wild-type or complemented strain ΔΔalp2F/alp2F-2G.
